# Morphometrics of a wild Asian elephant exhibiting disproportionate dwarfism

**DOI:** 10.1101/001594

**Authors:** Shermin de Silva, U. S. Weerathunga, T. V. Kumara

## Abstract

Dwarfism is a condition characterized by shorter stature, at times accompanied by differential skeletal growth proportions relative to the species-typical physical conformation. Causes vary and well-documented in humans as well as certain mammalian species in captive or laboratory conditions, but rarely observed in the wild. Here we report on a single case of apparent dwarfism in a free-ranging adult male Asian elephant (*Elephas maximus*) in Sri Lanka, comparing physical dimensions to those of other males in the same population, males in other populations, and records in previous literature. The subject was found to have a shoulder height of approximately 195cm, is shorter than the average height of typical mature males, with a body length of 218cm. This ratio of body length to height deviates from what is typically observed, which is approximately 1:1. In absolute height the subject was similar to the attributes of a captive elephant documented in 1955 in Sri Lanka, also said to be a dwarf, however the two specimens differed in the relative proportions of height vs. body length. The subject also exhibits a slight elongation of the skull. We discuss how this phenotype compares to cases of dwarfism in other non-human animals.

## Introduction

The general attribute shared across individuals exhibiting the various forms of dwarfism is reduced stature, though this alone is not diagnostic of dwarfism. Dwarfism may be divided into two types: proportionate, whereby individuals exhibit growth disturbance while maintaining isometric proportions relative to typical, and disproportionate, whereby physical proportions are scaled isometrically. There seems to be little agreement on the scientific criteria, phenotypic attributes, or physiological mechanisms which define proportionate dwarfism in humans, though these terms are applied to individuals from domesticated species such as cattle [1,2]. Disproportionate dwarfism (variously classified as achondroplasia, chondrodysplasia or diastrophic dysplasia) on the other hand, involves both shorter stature as well as changes to the proportions of the limbs, head and torso [3]. In humans it arises most commonly due to a single amino acid mutation in the protein known as fibroblast growth factor receptor 3 (FGFR-3), which can limit bone growth [3–5]. However, dwarfism can manifest from several distinct physiological mechanisms, only some of which have identified and heritable genetic components.

Dwarfism in non-human animals in the wild has rarely if at ever been documented. Among elephants in captivity, dwarfism has been recorded in at least two cases, one male and one female, shown in Figure 1 [6]. Anecdotal historic accounts exist of apparent dwarfism occurring in the wild in various parts of this species’ range [6], however these are not substantiated. Here we report on the case of dwarfism exhibited by a fully-grown male Asian elephant (*E. maximus maximus*) in Uda Walwe National Park, Sri Lanka [7] in comparison to individuals exhibiting typical growth habits and discuss this phenotype in relation to documented forms of dwarfism.

**Fig. 1.**
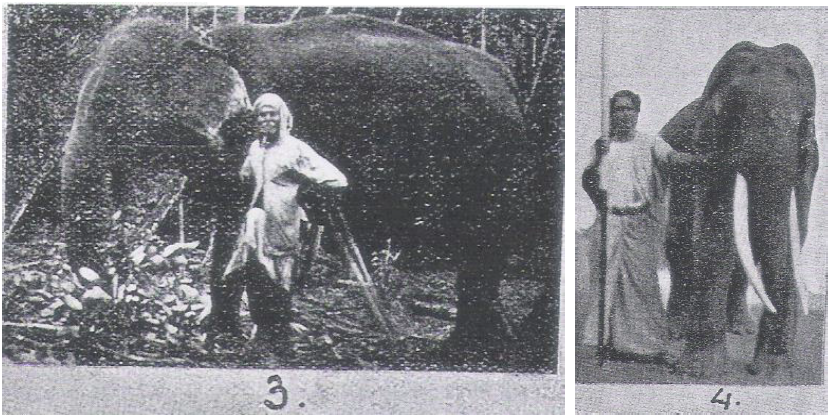
"Dwarf" elephants documented in captivity. Reproduced from [6], plate 18. The photograph marked (3) is of a 35 year old female described as a dwarf, owned by a person in Sri Lanka. The photograph marked (4) is of a 21 year old male captured in Mannar (north-west Sri Lanka), also described as a dwarf, with some physical measurements provided (see Table 1).

**Tab. 1.** Relative proportions of dwarf and wild type adult male Asian elephants in Sri Lanka. M459 (left) proposed dwarf, compared to M054 (right) a typical mature adult male. Numbers indicate ratios given in Table 1 (see methods) as follows: 1) Head to foot; 2) Width between the eyes; 3) Body length from base of tale to ear canal; 4) Shoulder height; 5) Foreleg length; 6) Top of skull to chin; 7) Top of skull to eye; 8) Eye to tusk sheath

|  | Anterior view |  | Lateral view |  |  |  |  |  |  |
| --- | --- | --- | --- | --- | --- | --- | --- | --- | --- |
|  | Head to foot | Eye width | Body length | Shoulder height | Foreleg | Head | Top of skull to eye | Eye to lip | Height (cm) |
| M054 | 11 | 2.8 | 14.5 | 14 | 8 | 6.5 | 4 | 3 | - |
| M343 | - | - | 13 | 13 | 7 | 5 | 3 | 2 | - |
| M034 | 11 | 2.6 | - | 12 | 6.5 | 5 | 3 | 2 | ~224 |
| M002 | 11 | 2.5 | 11.5 | 11.5 | 6.2 | 5 | 3 | 2 | - |
| M459 | 10 | 3 | 14 | 12.5 | 6 | 6.5 | 4 | 3 | ~152 |
| 21-yr old tusker | 12.7 | 3 | 13 | 12 |  |  |  |  | 183 |

## Study Site and Methods

Uda Walawe National Park (UWNP) is located in south-central Sri Lanka (latitudes 6°25’-6°34’N / 80°46’-81°00’E). Vehicle-based observations were made during long-term monitoring of this population conducted since 2006. The elephant population occupying UWNP numbers between 800-1200 [8], with a high degree of seasonality in use. In particular, males are known to range beyond this protected area, which is partially encircled by electric fencing. All individuals are photographed upon encounter, and individual-identification files based on natural features are maintained.

Shoulder height measurements were made opportunistically using one of two methods. The footprint circumference for a young male exemplifying average stature in this population was measured and the height was calculated as twice this circumference. This ratio appears to remain constant throughout an individual’s life [9]. The height of the subject was measured by proxy using a tree against which he had been standing.

In order to compare the subject’s physical stature to those of typical bulls, we assessed these proportions using photographs of known individuals (Figure 2). We used two views, anterior and lateral. The anterior view was used to determine a single ratio: height from top of head to toes vs. width of the distance between the eyes. The lateral view was used to calculate multiple ratios from the same photograph: the distance from the top of the skull to the eye, the eye to the base of the upper lip (the point where tusks or tushes would protrude), the length of the head, the length of the foreleg, shoulder height and back length measured from ear canal to the base of the tail. Only photographs in which subjects were directly facing or perpendicular to the angle of view of observers were used to limit angular distortions in measurements. Distortions from changes in elevation were minimized by always taking photographs from the same vehicle-based vantage point. Note that these provide relative rather than absolute morphometric attributes for comparative purposes.

## Results

The subject was first observed on July 11th 2013, video recorded, and assigned the ID M459 (Figure 2). M459 had not been identified in preceding years and therefore appeared to be a new arrival to UWNP. Temporal secretions indicated he was in the physiological state known as ‘musth,’ which is a period of heightened sexual activity and aggression [10]. The subject exhibited no unusual behavior but was un-habituated to observers and appeared fearful of humans.

**Fig. 2.**
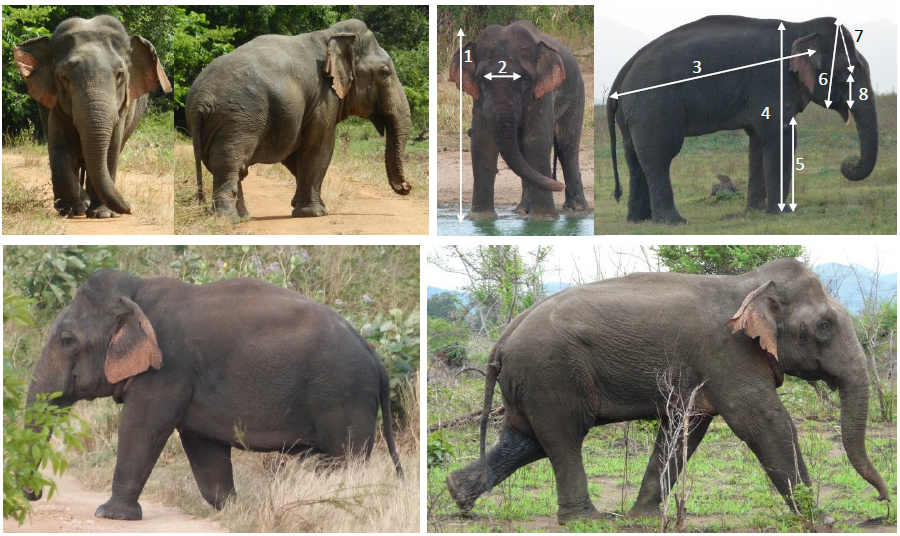
Dwarf M429 and wild type adult male M054

Morphometric details are provided in Table 1. The head is large relative to the length of the foreleg and the forehead is slightly elongated. M459’s estimated shoulder height was approximately 152cm. By comparison M343, a young adult male in this population, had already attained a shoulder height of 224cm (fore and hind foot circumference both equal to 112cm) and older, larger males are observed, such as M054. The subject is also well below the asymptotic height (∽250cm, range: 220-270cm) of Asian elephants measured elsewhere in the wild and captivity [9]. M459 exhibits shorter limbs, and slightly lengthened back relative to height compared to other mature males in this population. Since the young tusker captured in Sri Lanka in 1933 that was also documented as a dwarf (Figure 1, Table 1) was also wild-caught, it would have to be the first known case of dwarfism if it was indeed such. However, the proportions of this individual which was stated to have a shoulder height of 183cm and a back length of 198cm (measured from behind the ear, than from the ear canal, to the base of the tail), are more similar to wild type individuals despite the shorter absolute height.

## Discussion

We document an adult male individual occurring in a wild population of Asian elephants in Sri Lanka (*E. m. maximus*), M459, that exhibited isometric differences in physical dimensions relative to other adult males. These include shorter limbs, lengthening of the body, and possibly with macrocephaly, or enlargement of the skull. These attributes are consistent with what appears to be a case of disproportionate dwarfism. This individual moreoever exhibited obvious signs of sexual maturity, most notably temporal secretions indicative of musth. An interesting account of his confrontational behavior toward other males has also been recorded [7], evidencing apparently normal sexual activity. However, mating is generally preceded by intense competition, in which females actively run away from potential suitors, sometimes over multiple days. It is not known whether M459 could chase, guard, and physically mount a female in oestrus.

### Forms of size reduction across taxa

The term ‘dwarf’ and related term ‘pygmy’ do not designate particular conditions thus disambiguation is necessary. ‘Insular dwarfism’ refers to the phenomenon by which terrestrial island species exhibit reduction in size in comparison to mainland sister species through some evolutionary process [11,12]. The pygmy hippopotamus for instance (*Choeropsis liberiensis*) exemplifies this pattern as well as members of the Proboscidea occurring during the Pleistocene [11]. Mature bulls among the so-called pygmy elephants found on the island of Borneo (*E. m. borneensis*) can attain a shoulder height of at least 260cm (N. Othman, unpublished data) and thus are not necessarily smaller than their counterparts elsewhere in Asia, although more data are needed on their size distribution relative to other populations. Nevertheless, such a population-level trait is not what is described here, though Sri Lanka is an island.

### Disproportionate dwarfism in non-human animals

At this time, we are not aware of other documented cases of disproportionate dwarfism in the wild, although such accounts may exist. Among non-domesticated species, instances include individuals from two species of tamarins (*Sanguinus oedipus* and *Sanguinus fuscicollis*) resulting from matings between siblings, both of which were either stillborn or died as neonates [13]. These cases occurred in captivity however.

Instances of disproportionate dwarfism are widely documented among domestic and laboratory-reared animals and can arise from a variety of conditions with hormonal and genetic bases. Animal models such as *Mus* (mouse) and *Xenopus* (frog) have been used to investigate disruptions to the FGFR-3 pathway as implicated in humans, both at the genetic and protein levels [5]. Alternatively, mice can exhibit mutations in natriuretic peptide receptor 2 (Npr2) that also lead to achondroplastic characters. The former is an autosomal dominant gain-of-function mutation [4] while the latter is an autosomal recessive trait resulting from a loss-of-function mutation [14,15]. Both limit bone development, but in effect do so through opposing mechanisms either by enhancing the activity of a protein that limits bone growth (FGFR-3 disruption) or by inhibiting the activity of a protein that promotes bone growth (Npr2 disruption). Another candidate gene includes COL11A2, involved with the encoding of collagen, mutations in which are documented in mice and canids to give rise to shorter-limbed phenotypes due to abnormalities in cartilage development (chondrodysplasia) and may also be associated with hearing loss [16].

Disproportionate dwarfism occurs in larger-bodied domesticated ungulates such as cattle and horses as well. So-called "bulldog dwarfism" in Dexter cattle results from a homozygous lethal mutation in ACAN, a gene encoding Aggrecan, which is an essential component of cartilage [17]. While homozygotes incur severe deformities and typically abort midway through gestation, heterozygotes may exhibit milder dwarfism. Friesian horses exhibit a form of dwarfism in which both the fore and hind limbs are approximately 25% shorter relative to the breed’s typical conformation, with relatively larger heads, longer hooves and hyperextension of the fetlocks [18]. The head proportions themselves however appear relatively normal, unlike the effects observed with disruptions to FGFR1 or FGFR2, two candidate genes found within a region of the genome that appears to be highly associated with the dwarf phenotype in this breed [19]. Nor do the documented cases resemble phenotypes resulting from disturbance to PROP1, another candidate gene within the same region, that regulates growth hormone production. Thus the causal mechanism remains an open question. As growth disturbances can clearly arise from a wide array of genetic influences, the causes of the dwarfism exhibited by M459 cannot be verified at this time. The paucity of documented cases of disproportionate dwarfism in the wild may be due to reduced fitness and survival as a consequence of one or more genetic abnormalities, or a lack of outlets to report such observations outside medical literature.

## Acknowledgements

We would like to thank the warden of UWNP Mr. Pathirana, and personnel of the Department of Wildlife Conservation Sri Lanka for facilitating the ongoing research project. We also wish to thank Lim Tze Tshen for tracking down historical literature cited in this report as well as Nurzhafarina Othman for providing height measurements of Bornean elephants. A video clip of M459 may be viewed at http://youtu.be/ji8A6SoPaJI.

